# A role for the autophagic receptor, SQSTM1/p62, in trafficking NF-κB/RelA to nucleolar aggresomes

**DOI:** 10.1101/2020.04.09.034975

**Authors:** Ian T Lobb, Pierre Morin, Kirsty Martin, Xhordi Lieshi, Karl Olsen, Rory R Duncan, Lesley A Stark

**Author notes:** Joint first author (Ian T Lobb, Pierre Morin). Corresponding author: Lesley A Stark.

## Abstract

Elevated NF-κB activity is a contributory factor in many haematological and solid malignancies. Nucleolar sequestration of NF-κB/RelA represses this elevated activity and mediates apoptosis of cancer cells. Here we set out to understand the mechanisms that control the nuclear/nucleolar distribution of RelA and other regulatory proteins, so that agents can be developed that specifically target these proteins to the organelle. We demonstrate that RelA accumulates in intra-nucleolar aggresomes in response to specific stresses. We also demonstrate that the autophagy receptor, SQSTM1/p62, accumulates alongside RelA in these nucleolar aggresomes. This accumulation is not a consequence of inhibited autophagy. Indeed, our data suggest nucleolar and autophagosomal accumulation of p62 are in active competition. We identify a conserved motif at the N-terminus of p62 that is essential for nucleoplasmic-to nucleolar transport of the protein. Furthermore, using a dominant negative mutant deleted for this nucleolar localisation signal (NoLS), we demonstrate a role for p62 in trafficking RelA and other aggresome-related proteins to nucleoli. Together, these data identify a novel role for p62 in trafficking nuclear proteins to nucleolar aggresomes under conditions of cell stress, thus maintaining nuclear proteostasis. They also provide invaluable information on the mechanisms that regulate the nuclear/nucleolar distribution of RelA that could be exploited for therapeutic purpose.

**Significance:** Aberrant NF-κB activity drives many of the hallmarks of cancer and plays a key role in cancer progression. Nucleolar sequestration of NF-κB/RelA is one mechanism that switches off this activity and induces the death of cancer cells. Here we define a novel role for the autophagy receptor, SQSTM1/p62 in transport of nucleoplasmic NF-κB/RelA to nucleoli. Identification of this new trafficking mechanism opens up avenues for the development of a unique class of therapeutic agents that transport RelA and other cancer regulatory proteins to this organelle.

## Introduction

NF-κB is a family of essential transcription factors that play a key role in many of the hallmarks of cancer including cell proliferation, apoptosis, inflammation, angiogenesis, and metastasis (1-3). The most abundant form is a heterodimer of the RelA(p65)/P50 subunits. In most healthy cells, this heterodimer is retained in the cytoplasm by the inhibitor protein IκBα. However, in the majority of human malignancies, including solid and haematological tumours, NF-κB is aberrantly active in the nucleus which promotes cell growth, blocks apoptosis, perpetuates an inflammatory environment, accelerates disease progression and conveys resistance to therapy (4-6). Indeed, the paramount importance of NF-κB in all stages of tumorigenesis has led to its aggressive pursuit as a therapeutic target. However, most therapies developed to date target the main switch in NF-κB activation - cytoplasmic release from IκB – which causes systemic inhibition and severe toxicity.

Once in the nucleus, NF-κB proteins are modulated by a plethora of co-activators, repressors and post-transcriptional modifications (7-10). These nuclear regulatory pathways are also extremely important for cancer progression as they govern the genes that are activated or repressed in a specific context and hence, the downstream consequences on cell growth, death, differentiation and inflammation (11,12). However, unlike cytoplasmic activation of NF-κB, these post induction, nuclear mechanisms of regulation remain poorly understood. Further mechanistic understanding in this area is essential so that drugs can be developed that specifically target cells with aberrant nuclear NF-κB activity, exploiting cancer cell vulnerabilities.

One nuclear pathway us and others have identified to be important for inhibiting aberrant NF-κB transcriptional activity and inducing the death of cancer cells is nucleolar sequestration of RelA (13). In response to classic stimuli of NF-κB such as TNF, RelA is retained in the nucleoplasm, excluded from nucleoli (13). In contrast RelA translocates from the nucleoplasm to nucleoli in response to specific chemopreventatve agents (e.g aspirin, sulindac, sulindac suphone(13,14)), therapeutic agents (e.g proteasome inhibitors, the anti-tumour agent 2-methoxyestradiol (2ME2), TRK inhibition)(15,16)) and stress inducers (e.g UV-C radiation and serum starvation(13)). This differential distribution has important consequences as nucleoplasmic accumulation is associated with high NF-κB transcriptional activity and inhibition of apoptosis, while nucleolar sequestration is causally involved in repression of basal NF-κB transcriptional activity and the induction of apoptosis (13,15,16). Although some mechanisms that control this differential nuclear distribution have been reported, understanding is limited (17,18). Indeed, nucleolar trafficking and sequestration in general are poorly understood. Directing RelA and other regulatory proteins to nucleoli to eliminate diseased cells is a very attractive therapeutic option that can only be realised by further understanding in this area.

Here we demonstrate that RelA accumulates within intra-nucleolar aggresomes in response to therapeutic agents. We show that the nucleolar accumulation of RelA occurs alongside a flux of aggresome-related proteins into the organelle. We demonstrate that the autophagy receptor protein, SQSTM1/p62, also accumulates in nucleolar aggresomes in response these agents. Furthermore, we identify a nucleolar localisation signal at the N-terminus of p62 and, using a dominant negative mutant deleted for this motif, demonstrate a role for p62 in trafficking RelA and other aggresome-related proteins from the nucleoplasm to nucleoli. Together, these data identify a novel role for p62 in trafficking nuclear proteins to nucleolar aggresomes under conditions of stress. They also provide invaluable information on the mechanisms that regulate the nuclear/nucleolar distribution of RelA that could be exploited for therapeutic purpose.

## Materials and Methods

### Cells and reagents

SW480 cells were purchased from the American Type Culture Collections (ATCC) and maintained in L-15 medium (Gibco) supplemented with penicillin (100 IU/ml), streptomycin (100 ug/ml) and 10% fetal calf serum (FCS). Aspirin was supplied by Sigma (St. Louis, MS). It was solubilised in water using 10N NaOH, then the pH adjusted to 7.0. All other chemicals and reagents were from standard commercial source and of the highest quality.

### Immunocytochemical staining and image analysis

Immunocytochemistry was performed as previously described (13). Primary antibodies were: RelA (C20), RelA (F6) C23 (all Santa Cruz); Fibrillarin (Cytoskeleton); EIF4A1, EIF4G2, EIF4H (all Cell Signalling Technology); HSPA2 and 8 (Enzo Biosciences); SUMO-2/3 (Invitrogen); β-Catenin (BD Transduction Laboratories). Cells were mounted in Vectashield (Vector Laboratories) containing 1ug/ml DAPI. Images were captured using a Coolsnap HQ CCD camera (Photometrics Ltd, Tuscon, AZ, USA) Zeiss Axioplan II fluorescent microscope, 63 × Plan Neofluor objective, a 100 W Hg source (Carl Zeiss, Welwyn Garden City, UK) and Chroma 83 000 triple band pass filter set (Chroma Technology, Bellows Falls, UT, USA). Image capture was performed using scripts written for iVision 3.6 or Micromanager (https://open-imaging.com/). For each experiment, a constant exposure time was used. Image quantification was carried out using DAPI as a nuclear marker and C23 as a nucleolar marker along with ImageJ, image analysis software. At least 150 cells from at least 5 random fields of view were quantified for three independent experiments, or as specified in the text.

### Plasmids, SiRNAs and transfections

The GFP-RelA expression construct was a kind gift from E. Qwarnstrom (University of Sheffield) (19). DsRed-RelA was generated by subcloning RelA from GFP-RelA to pDsRed-C1 (Clonetech). GFP-Fibrillarin was a kind gift from A. Lamond (University of Dundee). Wild type GFP-p62 was a kind gift from Terje Johansen (20). GFP-p62ΔPB1 was generated by cloning cDNA from SW480 cells corresponding to isoform 2 of p62 [Δ1-84] into pEGFP-C1 (Clonetech). Full sequence analysis confirmed the sequence of p62ΔPB1 was identical to wild type protein, other than lack of the first 84aa. GFP-p62ΔPB1ΔNoLS was generated by sub-cloning aa122-440 from GFP-p62ΔPB1 into pEGFP-C1 while GFP-p62ΔPB1ΔUBA was generated by sub-cloning aa84-395 from GFP-p62ΔPB1 into this plasmid. The D_335_A/D_336_A/D_337_A mutations in the LC3 interacting domain were generated using a two-step approach. Firstly, the N and C termini of GFP-p62ΔPB1 were amplified using p62sens3/D335Arev and D335Afor/p62antisens primers. The PCR products were then digested with NotI (this restriction site is introduced by the mutation) and ligated together. After a second round of PCR with primers p62sens3/p62antisens the fragment was cloned into pEGFP-C1. All constructs were verified by sequencing. Plasmids were transfected into SW480 cells using Lipofectin (Gibco) as per manufacturer’s instructions. COMMD1 siRNA was generated and transfected into cells as described previously (17).

### Fluorescent correlation Spectroscopy (FCS): acquisition and analysis

SW480 cells were transfected with GFP-Fibrillarin, DsRed-RelA or control plasmids (pEGFP, pDsRed) then treated with aspirin (5mM, 16h). Hoescht 33342 (Sigma Aldrich) dye was added 30 min prior to imaging, to allow visualisation of DNA and nucleoli (unstained area in nucleus). All FCS recordings were acquired using a Leica SP5 SMD confocal microscope using a × 63 1.2NA HCX PL Apo water lens and 488 or 561 nm CW lasers. Photon fluctuation data routed through a Picoquant PRT 400 router were acquired at microsecond rates using external Single Photon Avalanche Photodiodes (MicroPhoton Devices, Italy). Autocorrelation traces were generated from the photon-counting histograms for each 5 to 30 s measurement using SymPhoTime v5.4.4 software (Picoquant, Germany). *In vitro* calibration traces were fitted using the Triplet model (three-dimensional free diffusion model with triplet state) with informed diffusion values to yield *V*_eff_ and *κ* values. Diffusion within the cells is expected to be anomalous; therefore, the anomaly parameter was not fixed to one. The anomaly parameter (*α*) measures the departure from free Brownian diffusion (*α*=1) to either superdiffusion (*α*>1) or subdiffusion (α<1) for a diffusing species. Autocorrelation curves with (*α*>1) display the sharpest decay, whereas the those with *α*<1 decrease quite slowly(21).

### Quantitative proteomics and bioinformatic analysis

Details of methods used to generate proteomic datasets are outlined in supplemental Fig. 3. The web-based tool, GOrilla (target v background), was used to determine whether proteins that increase in nucleoli in response to aspirin (>Log^2^ 0.5) are enriched for specific Gene Ontology (GO) terms. Background was set as all proteins isolated from nucleoli.

### Western Blot analysis and immunoprecipitations

Whole cell, cytoplasmic, nuclear and nucleolar extracts were prepared as previously described (13). Bradford assays (Bio-Rad) were used to measure protein content. Extracts (30 μg) were resolved using 5-12% sodium dodecyl sulfate-polyacrylamide gels, and immunoblotting performed by standard procedures. Primary antibodies used were: RelA (C20), C23, GFP, B23/nucleophosmin, RPA194 (all from santa Cruz), p62 (BD biosciences), α–tubulin (SIGMA), α-actin (SIGMA), SUMO2/3 (Invitrogen), β-catenin (BD Transduction Laboratories), ubiquitin (Dako). α-Tubulin (Sigma).Immunoprecipitations were performed on whole cell lysates from aspirin treated cells as previously described (17)using anti-RelA antibodies (C20, Santa Cruz) followed by p62 (anti-Mouse, BD Biosciences) immunoblots.

## Results

### Fluorescent correlation spectroscopy (FCS) reveals RelA accumulates in nucleolar aggregates

Proteins that translocate into nucleoli generally do not distribute evenly throughout the organelle, but accumulate in nucleolar bodies. These bodies have been independently termed nucleolar aggresomes, intra-nucleolar bodies, detention centres or cavities (22-27). To help understand the nuclear distribution of RelA mechanistically, we determined whether the protein resides within such aggregates.

Firstly, we used immunocytochemistry to examine the sub-nucleolar localisation of RelA in response to aspirin and MG132. Aspirin was used as a tool compound as we are interested in the mechanisms underlying its anti-tumour activity (28). We found, as has previously been described, that exposure to these agents induces an increase in size of nucleoli and segregation of nucleolar marker proteins to the periphery of the organelle (Figs. 1A and B)(29). We also found that RelA accumulates in very distinct foci in the centre of these enlarged nucleoli (Figs. 1A and B). Both endogenous and GFP-tagged RelA formed nucleolar foci, suggesting this is a genuine sub-nucleolar localisation pattern (Figs. 1A-C).

**Figure 1:**
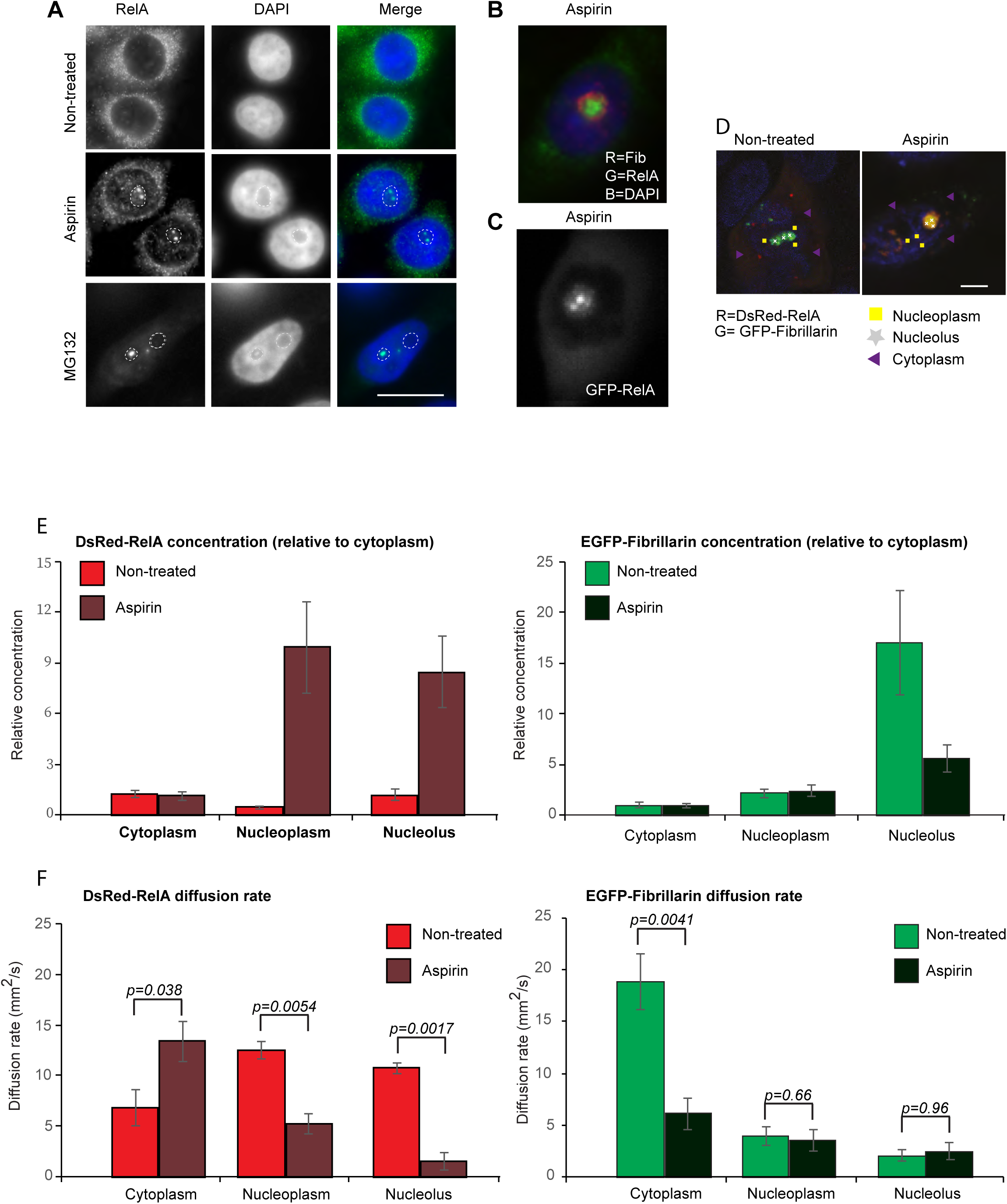
RelA accumulates in nucleolar aggregates, as determined by fluorescent correlation spectroscopy (FCS) **A** Immunomicrographs (X63) showing RelA foci within enlarged nucleoli (areas devoid of DAPI staining) in control (non-treated), aspirin (3mM, 16h) or MG132 (25uM, 8h) treated SW480 cells. Nuclei are stained with DAPI. Scale bar 10um. **B** SW480 cells were treated with aspirin as in A. Immunomicrograph depicts nodular RelA staining in the centre of enlarged, segregated (as evidenced by peripheral fibrillarin (Fib)) nucleoli. R=red. G=green, B=blue. **C** SW480 cells were transfected with GFP-tagged RelA 24h prior to treatment with aspirin as in A. The sub cellular distribution of GFP-RelA was determined in live/adherent cells. **D-F** SW480 cells were transfected with Dsred-RelA and GFP-Fibrillarin, treated with aspirin (0-5mM 16h) then analysed by FCS. **D** Representative confocal images of SW480 cells co-expressing DsRed-RelA (R=red) and EGFP-Fibrillarin (G=green). Hoescht 33342 (blue) staining of the nucleus allowed identification of (Hoescht-free) nuclolei. FCS recordings were made from spots in cytosolic (purple triangles), nucleoplasmic (yellow squares) and nucleolar (white crosses) regions of the same cells. **E** The concentration of each labelled protein was measured by FCS in each of the three compartments in control and aspirin treated cells. Concentrations in the nucleoplasm and nucleolus are expressed relative to the cytosolic concentration of the same cell, to account for differences in expression levels. Error bars represent the SEM of n=18 cells, over 2 biological repeats. **F** Diffusion rate measurements for DsRed-RelA (left) and EGFP-Fibrillarin (right) in each of the three compartments in control and aspirin-treated cells. The average of the two experiments (n=9 for each) is shown. Error bars represent the SEM and p-values are derived using the Kruskal-Wallis Test. See also Supplemental Figures 1 and 2 for additional controls.

To further explore the nature of nucleolar RelA foci, we utilised fluorescent correlation spectroscopy (FCS). This approach is ideal as it can report molecular concentration and diffusion rate at multiple points within a cell at a microsecond timescale (30). Processes that slow down the diffusion rate of molecules, such as binding in large complexes or aggregates, are visible by a shift of the FCS curve to longer lag times.

SW480 cells were transiently transfected with DsRed-RelA and EGFP-fibrillarin, exposed to aspirin, then FCS employed to quantify the concentration and diffusion rates of the tagged proteins within the cytoplasm, nucleoplasm and nucleoli (Fig. 1D). Supplemental Fig1 shows the average, normalised experimental and fitted autocorrelation curves for DsRed-RelA in each compartment, with and without aspirin treatment. The mean t1 is marked. Examples of individual recordings from each compartment are also shown (Supplemental Fig 1).

Analysis of FCS curves revealed that aspirin induces a significant increase in the concentration and decrease in the diffusion rate of DsRed-RelA in nucleoli, in keeping with accumulation of the protein within aggregates in this compartment (Figs 1E and F). A reduction in the diffusion rate of DsRed-RelA was also observed in the nucleoplasm, although to a slightly lesser degree. In contrast, the diffusion rate of DsRed-RelA increased in the cytoplasm in response to the agent (Figs 1E and F), consistent with release of RelA from the IκB inhibitor complex. Unlike DsRed-RelA, the diffusion rate of EGFP-fibrillarin remained constant in the nucleoplasm and nucleoli (Fig 1F), as did DsRed and EGFP controls (Supplemental Fig 2). The nucleolar concentration of EGFP-fibrillarin appeared to decrease in response to aspirin. However, this can probably be explained by segregation of this protein to the nucleolar periphery, out with the location of the data collection points (Figs.1B, D, and E). Interestingly, in contrast to DsRed-RelA, aspirin induced a significant reduction in the cytoplasmic diffusion rate of EGFP-fibrillarin (Fig. 1F). We currently have no explanation for this result. However, it does confirm that the agent has compartment and protein specific effects on protein mobility.

Together, these data establish the power of FCS for investigating NF-κB signalling in distinct cellular compartments. They also provide convincing evidence that RelA resides within aggregates in nucleoli following exposure to specific therapeutic agents.

### RelA accumulates in nucleoli alongside heat shock factors, translation factors and proteins of the ubiquitin proteasome system

In a related study, Stable Isotope Labelling of Amino acids in Culture (SILAC)-based quantitative proteomics was used to investigate aspirin- and MG132-induced changes to the cytoplasmic, nucleoplasmic and nucleolar proteomes (Supplemental Fig. 3). To explore the global proteomic landscape in which RelA aggregates are observed, we interrogated these datasets. Particularly, those derived from 10h aspirin (N=3) and MG132 (N=2), as immunocytochemistry indicated nucleolar RelA is evident under these conditions (Fig. 2A).

**Figure 2.**
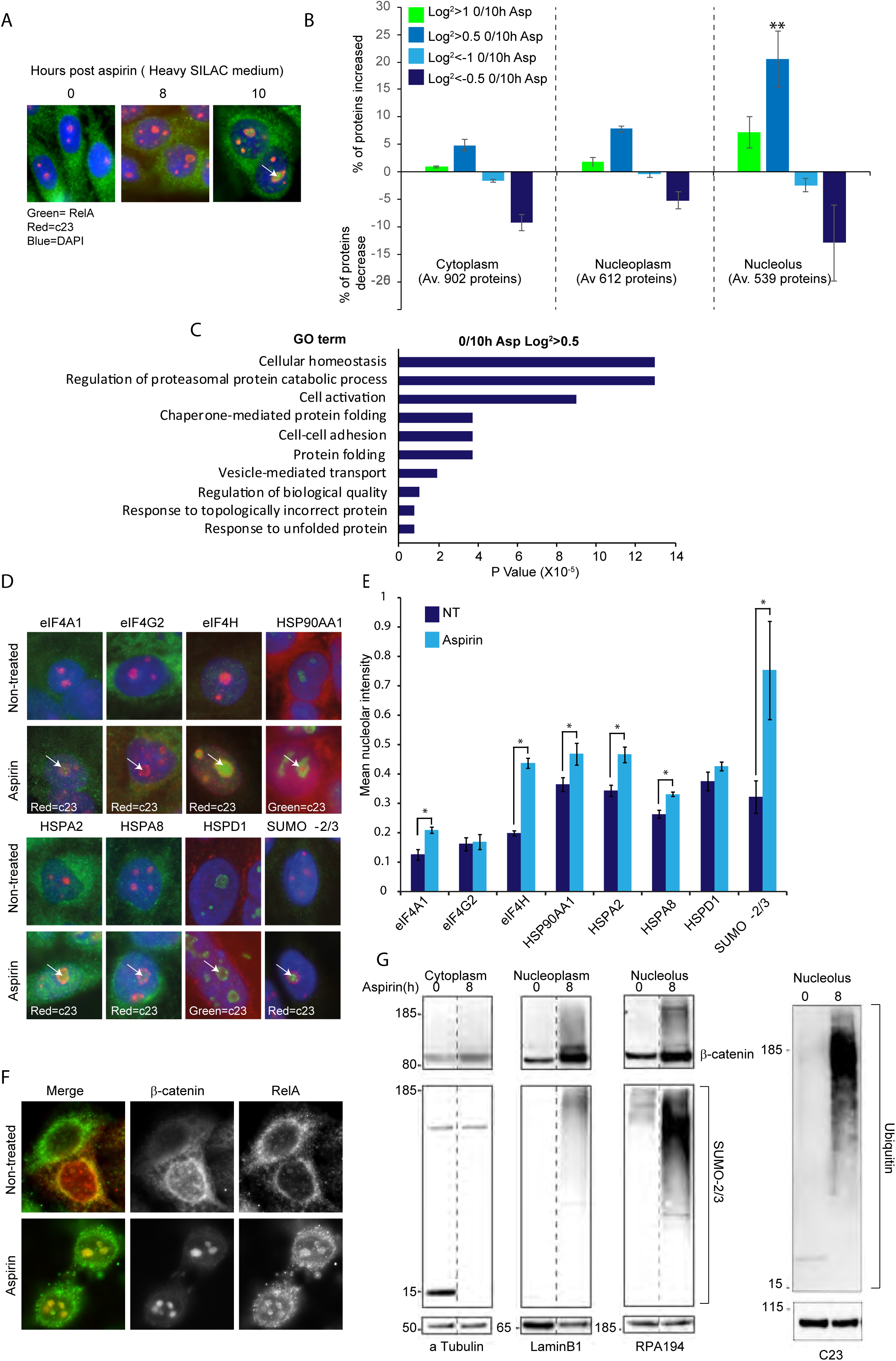
RelA accumulates in nucleoli alongside heat shock factors, translation factors and proteins of the ubiquitin proteasome system. **A** SW480 cells were exposed to aspirin (10mM) for 0-10hrs in heavy (Lys^4^Arg^8^) Stable Isotope Labelling of Amino Acids in Culture (SILAC) medium. Immunomicropgraphs show the localisation of RelA and C23 (nucleolar marker protein). DAPI (blue) marks nuclei. Arrow indicates nucleolar aggregates of RelA. **B** SW480 cells were grown in SILAC media then either left untreated or treated with aspirin (10mM) for 10h. Cells were pooled, cytoplasmic, nucleoplasmic and nucleolar fractions isolated, run on SDS page gels, analysed by mass spectrometry and peptides quantified using MaxQuant software (see Supplemental Figure 3 for full details). All data was plotted as normalized Log^2^ values against the control. Three biological repeats were performed (N=3). The graph depicts the average number of proteins that increased (Log^2^ 0/10h>0.5 or >1) or decreased (Log^2^ 0/10h<-0.5 or <-1) in each compartment (+/− SEM). The average (Av.) number of proteins isolated from each fraction is given. ** P=0.015 (2 tailed students ttest) comparing Log^2^ >0.5 cytoplasm to Log^2^ >0.5 nucleoli. **C** Proteins with a 0/10h aspirin Log2 ratio of >0.5 were subjected to gene ontology (GO) analysis. The top ten terms are shown. **D to F** SW480 cells were either non-treated or treated with aspirin (10mM, 8h) then immunocytochemistry performed with the indicated antibodies. **D** Immunomicrographs showing protein localisation. Arrows highlight nucleolar aggregates. C23 acts as a nucleolar marker. DAPI (blue) identifies nuclei. **E** ImageJ software was used to quantify the intensity of the proteins in nucleoli (as depicted by c23). A minimum of 200 nucleoli were analysed per experiment from 10 fields of view. The mean (+/−SEM) is shown. N=3. **F** Immunomicrographs showing co-localisation of RelA and β-catenin in nucleoli in response to aspirin. **G** SW480 cells were treated with aspirin as indicated, cells fractionated into cytoplasmic, nucleoplasmic and nucleolar fractions, then western blot analysis performed with the indicated antibodies. α-tubulin (cytoplasm), laminB1 (nucleoplasm) and RPA194/c23 (nucleoli) act as loading controls.

Our analysis revealed that both aspirin and MG132 have a particularly significant effect on the nucleolar proteome, compared to the cytoplasmic and nucleoplasmic proteomes (Fig. 2B and supplemental Fig. 3C). They also revealed both agents induce an influx of proteins into the organelle (Fig. 2B and supplemental Fig. 3C). Gene ontology analysis of proteins that increased in nucleoli in response to aspirin indicated enrichment (P<0.00001) for terms associated with regulation of biological quality, protein folding and response to topologically incorrect protein (Fig. 2C). Immunocytochemistry and western blot analysis were used to analyse specific proteins, and protein families, found to accumulate in nucleoli in response to aspirin. These data confirmed that aspirin induces nucleolar accumulation of translational regulators (eIF4A1, eIF4H), chaperones (HSP90AA1, HSPA2, HSPA8), UPS proteins (ubiquitin, SUMO-2/3) and cell adhesion molecules (β-catenin) (Figs. 2D to G). Furthermore, such proteins co-localised with RelA in aggregates in this compartment (Fig 2F). Interestingly, there was specificity in aspirin-mediated nucleolar accumulation of regulatory proteins as HSPA family chaperones accumulated in the compartment, but not HSPD1. Similarly, EIF4A1 and EIF4H translocated to nucleoli after aspirin treatment, but not EIF4G2. RelA was not identified in any compartment using SILAC based proteomics, possibly due to difficulties detecting RelA using LC-MS/MS.

One mechanism by which MG132 alters cell phenotype is the induction of proteotoxic stress. Given the types of proteins that accumulate in nucleoli in response to aspirin, and the similarity between aspirin and MG132 effects on the nucleolar proteome, we considered that aspirin also induces proteotoxic stress. A marker for this is accumulation of insoluble proteins and so, we measured the percentage of detergent insoluble protein following aspirin and MG132 exposure (Supplemental Fig. 3D). We found that aspirin did indeed cause an accumulation of insoluble protein in colon cancer cells. This occurred in a time dependent manner and was significant 5-8 hours following exposure to the agent, which precedes the induction of apoptosis (31). Furthermore, the increase in insoluble protein was comparable to that observed with MG132 (Supplemental Fig. 3D).

Together, these data provide compelling evidence that RelA accumulates in nucleolar aggregates alongside specific chaperones and targets of the ubiquitin proteasome pathway. It also confirms that proteotoxic stress has a particularly significant effect on the nucleolar proteome (24).

### Identification of p62 as a nucleolar shuttling protein

One protein of considerable interest with regard to the shuttling of RelA to nucleolar aggresomes is P62/Sequestosome 1 (SQSTM1). P62 is a multifunctional protein that acts as a signalling scaffold and autophagy cargo receptor(32,33). It colocalises with cytoplasmic aggregates and plays a significant role in their clearance (20). It also shuttles to the nucleus, although its role in this compartment is still not clear (34). It interacts with the autophagosomal marker, LC-3B I/II, which has previously been shown to be present in nucleoli {Beg, 1996 53 /id}. Furthermore, it is known to interact with RelA and reportedly transports RelA to autophagosomes for degradation (35). Hence, we considered that p62 may play a role in the formation or clearance of RelA containing nucleolar aggresomes.

To begin to address this question, we firstly determined whether, as described for other stimuli (35), p62 and RelA interact in response to aspirin. Indeed, immunoprecipitation assays demonstrated that the agent induces a transient interaction between the two proteins (Fig. 3A). Next, we utilised immunocytochemistry and western blot analysis to examine the sub cellular localisation of p62. These data revealed that aspirin induces the rapid nuclear accumulation of p62, followed by nucleolar translocation of the protein (Figs. 3B and C). Furthermore, nucleolar translocation of p62 occurred in parallel with nucleolar translocation of RelA, subsequent to their observed interaction (Figs. 3A and B). On a single cell basis, there was a direct correlation between cells that showed nucleolar RelA and those that showed nucleolar p62 (Supplemental Fig. 4A). In addition, p62 co-localised with RelA in foci in this compartment (Fig. 3D). We did not detect an interaction between p62 and RelA at the time point they were present in foci (8h), which may reflect the insoluble nature of the aggregates. P62 was also found to accumulate in nucleoli alongside RelA in response to MG132 (Figs. 4B and E). In contrast, LC-3B accumulated in the cytoplasm in response to aspirin (Fig. 3F). Together, these data indicate that aspirin induces the nuclear/nucleolar accumulation of p62 and suggest the intriguing possibility that this protein transports RelA to nucleolar aggregates. To test this possibility, we set out to examine the pathways underlying nucleolar translocation of p62.

**Figure 3.**
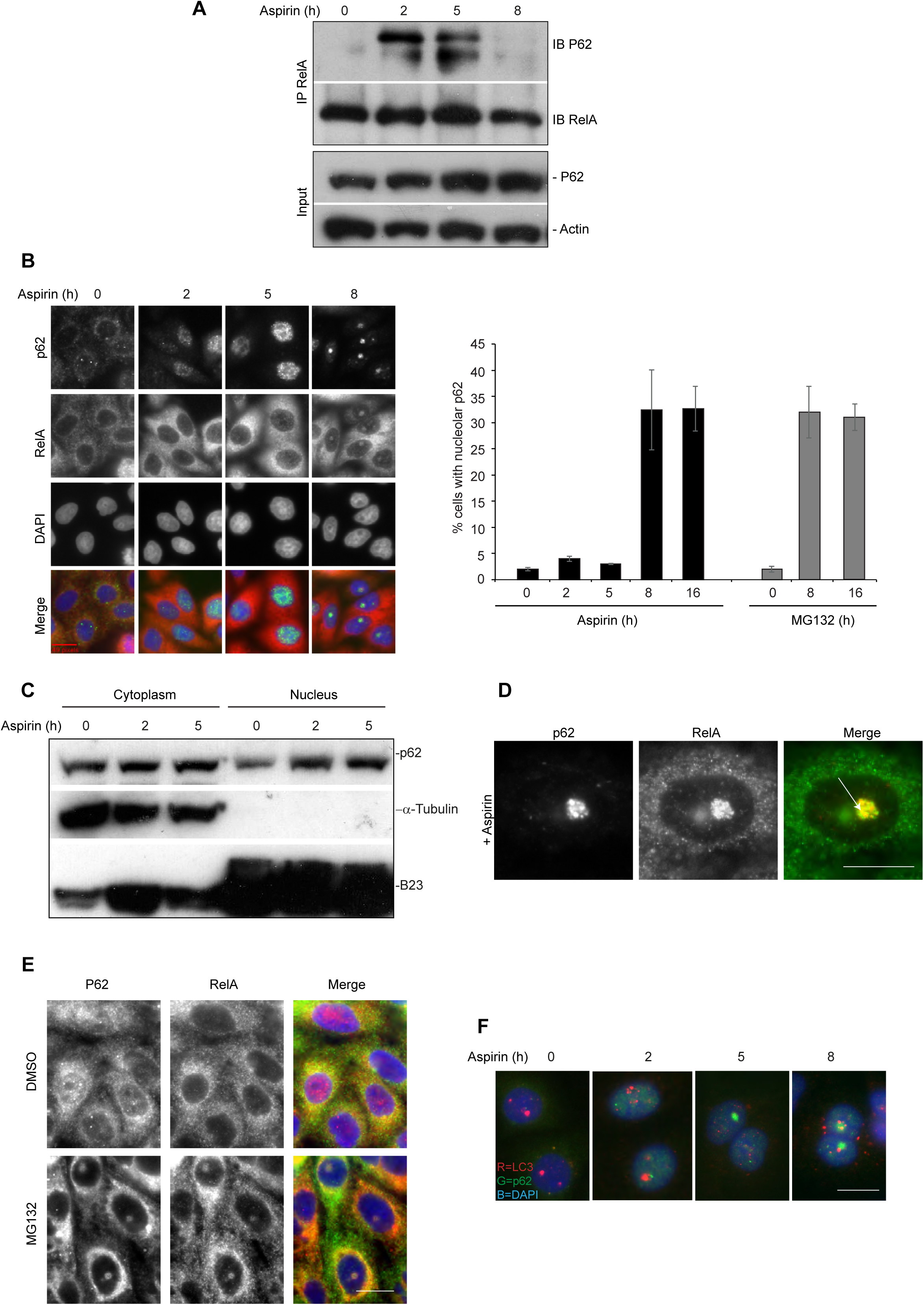
p62 shuttles to nucleoli in parallel with RelA. **A** RelA and p62 interact in response to aspirin. SW480 cells were treated with aspirin (10mM) for the times specified. Immunoprecipitations (IPs)were carried out on whole cell lysates using antibodies to RelA. Precipitated proteins were subjected to immunoblot (IB) analysis with antibodies to p62 and RelA. Input levels are shown. **B** Left: SW480 cells were treated with aspirin (10mM) for the times specified then immunocytochemistry performed with the indicated antibodies. DAPI depicts nuclei. Right:The percentage of cells showing nucleolar p62 was determined in at least 200 cells in 10 fields of view. Mean of three individual experiments is shown (+/−SEM). **C** Cells were treated with aspirin as in (B) then anti-p62 western blot analysis performed on cytoplasmic and nuclear fractions. α-tubulin (cytoplasm) and B23 (nuclei) act as loading controls. **D** RelA and p62 co-localise in nucleolar aggregates. Immunomicrograph (X63) showing the nucleolar localisation of RelA and p62 in aspirin (5mM, 16h) treated SW480 cells. **B and E** SW480 cells were treated with MG132 (25uM) for 0/16h. Immunocytochemistry was performed with antibodies to p62 and RelA. Nuclei are identified by DAPI stain (blue). Scale bars 10um throughout. **F** SW480 cell were treated with aspirin as in B then immunocytochemistry performed with antibodies to p62 and LC3B.

**Figure 4.**
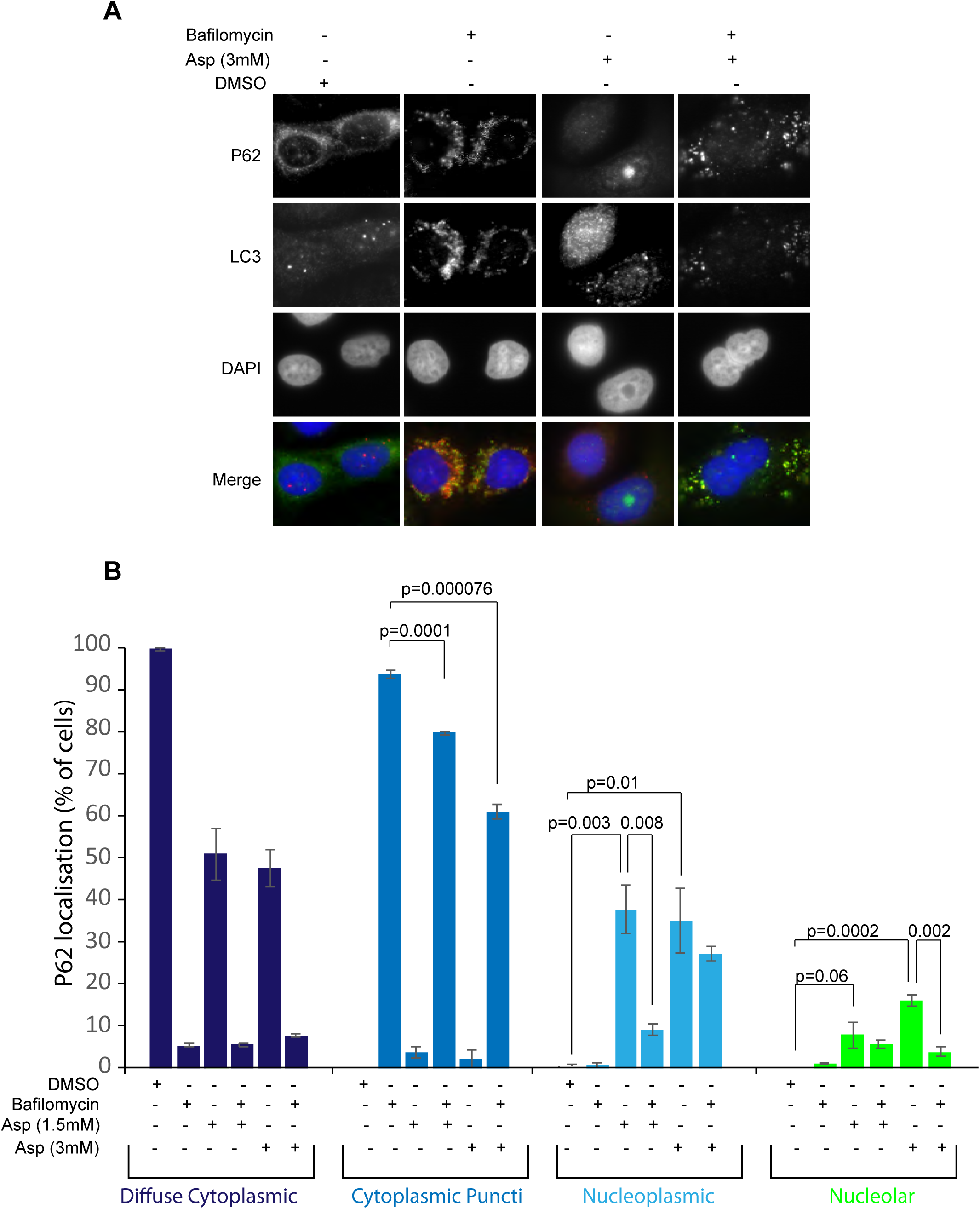
Interplay between autophagic and nucleolar accumulation of p62. **A and B** SW480 cells were pre-treated with DMSO or BafilomycinA1 (100nM) then exposed to aspirin (0-3mM, 16h). **A** Immunomicrographs (X63) show the localisation of p62 and the autophagosomal marker, LC-3. B The percentage of cells with the stated sub-cellular distribution of p62 was determined in at least 200 cells from 5 fields of view. The mean +/− SEM is shown. N=3. P values were derived using a two tailed students T test. Scale bars=10um.

### Autophagy and the formation of nucleolar aggresomes

Given the importance of autophagy in the clearance of p62 and its cargo proteins(32), we hypothesised that there may be interplay between autophagy and nuclear/nucleolar translocation of the protein. Specifically, we speculated that nucleolar aggregates form as a consequence of overwhelmed or blocked autophagy. Contrary to this suggestion, we found that specifically blocking the formation of autophagosomes, using the inhibitor of autophagasomal degradation, BafilomycinA1, did not induce nuclear/nucleolar accumulation of p62. Rather, as has been well documented previously, the protein accumulated in autophagosomes (LC3B positive cytoplasmic foci) in response to the agent (Figs. 4A and B).

Next we determined whether inhibiting autophagic degradation modulated aspirin-mediated nucleolar translocation of p62 (Figs. 4A and B). Quantitative immunocytochemistry confirmed that BafilomycinA1 induced cytoplasmic aggregation of p62 (Fig. 4B). However, the percentage of cells showing this localisation pattern was significantly reduced when cells were exposed to aspirin. Furthermore, this occurred in a manner dependent on aspirin dose (Fig. 4B). Similarly, when cells were exposed to aspirin alone there was a significant increase in those showing nucleoplasmic/nucleolar p62, but this was significantly reduced in the presence of BafilomycinA1 (Figs. 4A and B). Lysotracker, which detects active lysosomes, confirmed that BafilomycinA1 was equally active in the presence and absence of aspirin (data not shown). These data suggest that the pathways that target p62 to autophagosomes, and those responsible for nuclear/nucleolar translocation of the protein, may be in active competition.

### PB1, LIR and UBA domains are dispensible for nucleolar translocation of p62

To translocate to autophagosomes, p62 firstly has to homodimerize, which is dependent on the N-terminal PB1 domain (34). This domain also allows p62 to interact with the autophagy receptor NBR1 and the protein kinases ERK, MEKK3, MEK5, PKCζ, and PKCλ/ι (36). To further investigate the mechanisms underlying nucleolar translocation of p62, we utilised a GFP-tagged fusion protein deleted for this domain (GFP-p62ΔPB1) (Fig. 5A). GFP-P62 wild type (WT) and GFP-p62ΔPB1 were transfected into SW480 cells and the subcellular localisation of GFP-tagged proteins quantified in the presence and absence of aspirin (0, 3mM) (Figs 5B and C). In the absence of aspirin, GFP-p62WT was present in cytoplasmic puncti. In contrast GFP-P62ΔPB1 was diffuse within the cytoplasm, as would be expected given that it had lost the ability to self-aggregate. Following aspirin exposure, both proteins translocated into the nucleoplasm and accumulated in nucleoli. However, this effect is significantly more pronounced for GFP-P62ΔPB1 than for WT protein (Figs. 5B and C). These data indicate that the N-terminal PB1 domain is not required for nucleolar translocation of p62. The increased effect on GFP-P62ΔPB1 is also in keeping with our suggestion that there is competition between aggregation of p62 at autophagosomes and nucleolar targeting of the protein.

**Figure 5.**
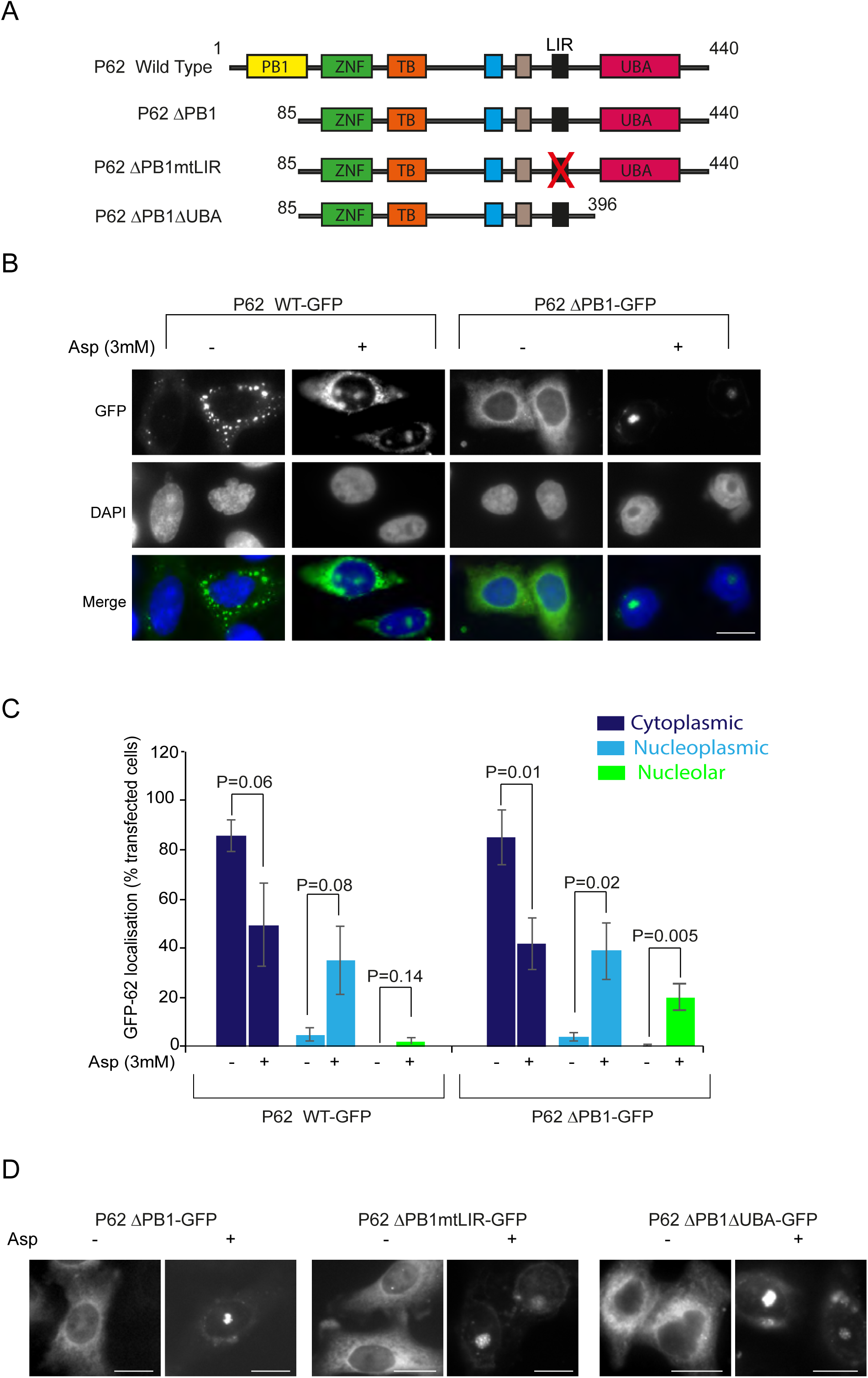
Nucleolar translocation of p62 does not require dimerization, LC3 or ubiquitin binding. **A** Schematic of p62 domain structure and the mutants generated. PB1:Phox and Bem1p, LIR:LC3 interacting region, ZNF:Zn Finger Domain, UBA:Ubiquitin-associated domain,. **B and C** SW480 cells were transfected with GFP-P62 Wild type (WT)- and P62ΔPB1 expression constructs then treated with aspirin (3mM, 16h). **B** Immunomicrographs (X63) of fixed cells. Nuclei are stained with DAPI. **C** The percentage of cells showing the indicated sub-cellular distribution of GFP-p62 (WT or ΔPB1) was determined in at least 100 transfected cells over more than 5 fields of view. Mean+/−SEM is shown. N=3. **D** SW480 cells were transfected with the indicated mutants and treated as above. Immunomicrographs (63X) of fixed cells are shown. Scale bars=10um.

Next, we explored the role of the LC3 interacting region (LIR) and the ubiquitin binding (UBA) domain in nucleolar transport of p62 (Fig. 5A). The UBA is required for p62 to bind to ubiquitinated proteins. LC3 binding, via the LIR, is essential for incorporation into autophagosomes. We generated an LC3 mutant (D_335_A/D_336_A/D_337_A) that decreases P62-LC3 binding by 75% (34) (GFP-P62ΔPB1mtLC3), and a C-terminal truncated version of the protein missing the UBA domain (GFP-P62ΔPB1ΔUBA). The mutants were generated in GFP-P62ΔPB1, as nucleolar translocation of this protein was not impeded by cytoplasmic aggregation. Analysis of fixed cells indicated that GFP-P62ΔPB1mtLC3 and GFP-P62ΔPB1ΔUBA accumulate in nucleoli in response to aspirin in a similar manner to GFP-P62ΔPB1 (Fig. 5D), suggesting these domains are also not essential for nucleolar targeting of the protein.

### Identification of a P62 nucleolar localisation signal

Nucleolar localisation of multiple proteins is dependent on the presence of a specific nucleolar localisation signal (NoLS)(37-39). To determine if p62 has such a signal, we submitted the sequence to an NoLS prediction program(40). Using this approach, we identified a domain of interest between aa 94 and 116 (IFRIYIKEKKECRRDHRPPCAQE) (Fig. 6A). This domain is located just after PB1, preceding the nuclear localisation and nuclear export signals (34). Alignment software revealed a stretch of basic amino acids within this region (aa 100 and 110, KEKKECRRDHR) that is common to known NoLSs (Fig. 6B). Bioinformatic analysis also revealed that this domain is conserved between species and is evident down to Danio Rerio, suggesting a strong selection pressure for the region (Fig. 6C).

**Figure 6.**
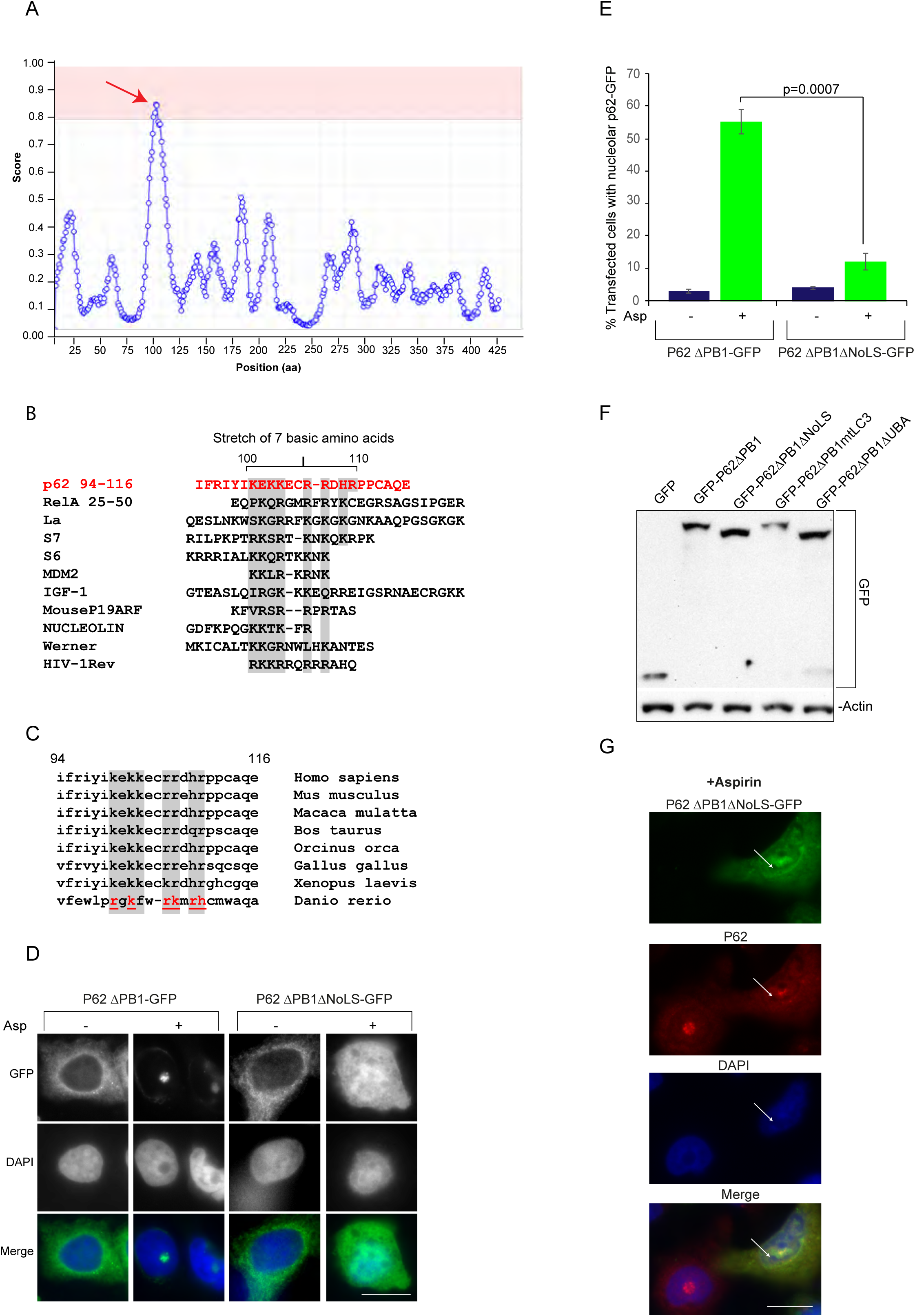
Identification of a p62 nucleolar localisation signal (NoLS). **A** The p62 sequence was submitted to NoD, a Nucleolar Localisation Signal (NoLS) prediction program {Scott, 2011 #1702}. The output, predicting an NoLS between aa 94 and 116 (score >0.8), is shown. Red arrow indicates a domain of interest **B** The predicted p62 NoLS [94-116] was aligned with NoLSs from other nucleolar shuttling proteins. Grey highlights a stretch of basic residues which is common for NoLSs. **C** aa 94-116 (putative NoLS) of Homo sapien p62 was aligned with p62 from the given species. The most divergent sequence (Danio rerio) retains 6 basic residues in common (in red and underlined). **D and E** SW480 cells were transfected with GFP-P62ΔPB1 or GFP-p62ΔPBΔNoLS then treated with aspirin as in Fig. 5. **D** Immunomicrographs (X63) of fixed cells showing the localisation of the GFP-tagged proteins. **E** The percentage of cells showing nucleolar GFP-p62 was determined in at least 100 transfected cells. The mean +/− SEM is shown. N=3. P values were derived using a two tailed students T test. **F** Anti-GFP-immunoblot performed on whole cell extracts from SW480 cells transfected with the specified constructs. Actin acts as a loading control. **G** GFP-p62ΔPBΔNoLS acts as a dominant negative mutant. SW480 cells were transfected with GFP-p62ΔPBΔNoLS then treated with aspirin as above. Immunocytochemistry was performed with primary antibodies to P62 and a TXred conjugated secondary antibody (X63). Nucleolar p62 can be observed in the non-transfected cell, which is blocked in the cell expressing the NoLS mutant (arrow indicates nucleoli as depicted by area devoid of DAPI). Nuclei are stained with DAPI throughout.

To determine the role of the putative NoLS in nucleolar translocation of p62, we further deleted GFP-p62ΔPB1 to create a construct that was deleted for residues 1-120 (GFP-p62ΔPB1ΔNoLS). GFP-p62ΔPB1ΔNoLS was transfected into SW480 cells and its localisation in the presence and absence of aspirin compared to GFP-p62ΔPB1. These studies revealed that both GFPp62ΔPB1 and GFP-p62ΔPB1ΔNoLS were diffuse in the cytoplasm prior to aspirin treatment and translocated from the cytoplasm to the nucleoplasm in response to the agent (Fig. 6D). However, while GFP-p62ΔPB1 accumulated in nucleoli, GFPp62ΔPB1ΔNoLS remained in the nucleoplasm (Fig. 6D). Image quantification confirmed that the significant increase in nucleolar GFP-p62ΔPB1 observed in response to aspirin was lost upon deletion of the putative NoLS (Fig. 6E). Western blot analysis confirmed that GFP-p62ΔPB1ΔNoLS is expressed at a similar level to GFP-p62ΔPB1 and other mutants, excluding differential expression level as an explanation for the differential localisation (Fig. 6F). These data show for the first time that p62 has a bona fide nucleolar localisation signal that is required for transport of the protein from the nucleoplasm to the nucleolus under conditions of stress.

Immunocytochemistry with antibodies to endogenous p62 revealed that expression of GFP-p62ΔPB1ΔNoLS blocks nucleolar translocation of endogenous protein (Fig. 6G), which is unexpected given that GFP-p62ΔPB1ΔNoLS does not have a self-binding domain. Nevertheless, it does indicate this mutant can be used as a tool to explore the role of p62 in nucleolar transport of cargo proteins.

### Nucleolar translocation of p62 is required for the formation of RelA nucleolar aggresomes

We next utilized the p62 ΔNoLS deletion mutant to directly test whether the nucleolar localization of p62 is causally involved in nucleolar translocation of RelA. Using RelA immunocytochemistry performed on transfected SW480 cells, we found, as expected, that cells expressing GFP-P62ΔPB1 showed nucleolar localization of both p62 and RelA in response to aspirin (Fig. 7A). However, in cultures expressing GFP-P62ΔPB1ΔNoLS, there was a significant reduction in transfected cells showing nucleolar RelA (Fig. 7B). Instead, both p62 and RelA remained in the nucleoplasm (Fig. 7A). In contrast, adjacent non-transfected cells showed aspirin-mediated nucleolar accumulation of both p62 and RelA (Fig. 7A). These results provide compelling evidence that p62 is required to transport RelA from the nucleoplasm to nucleoli.

**Figure 7.**
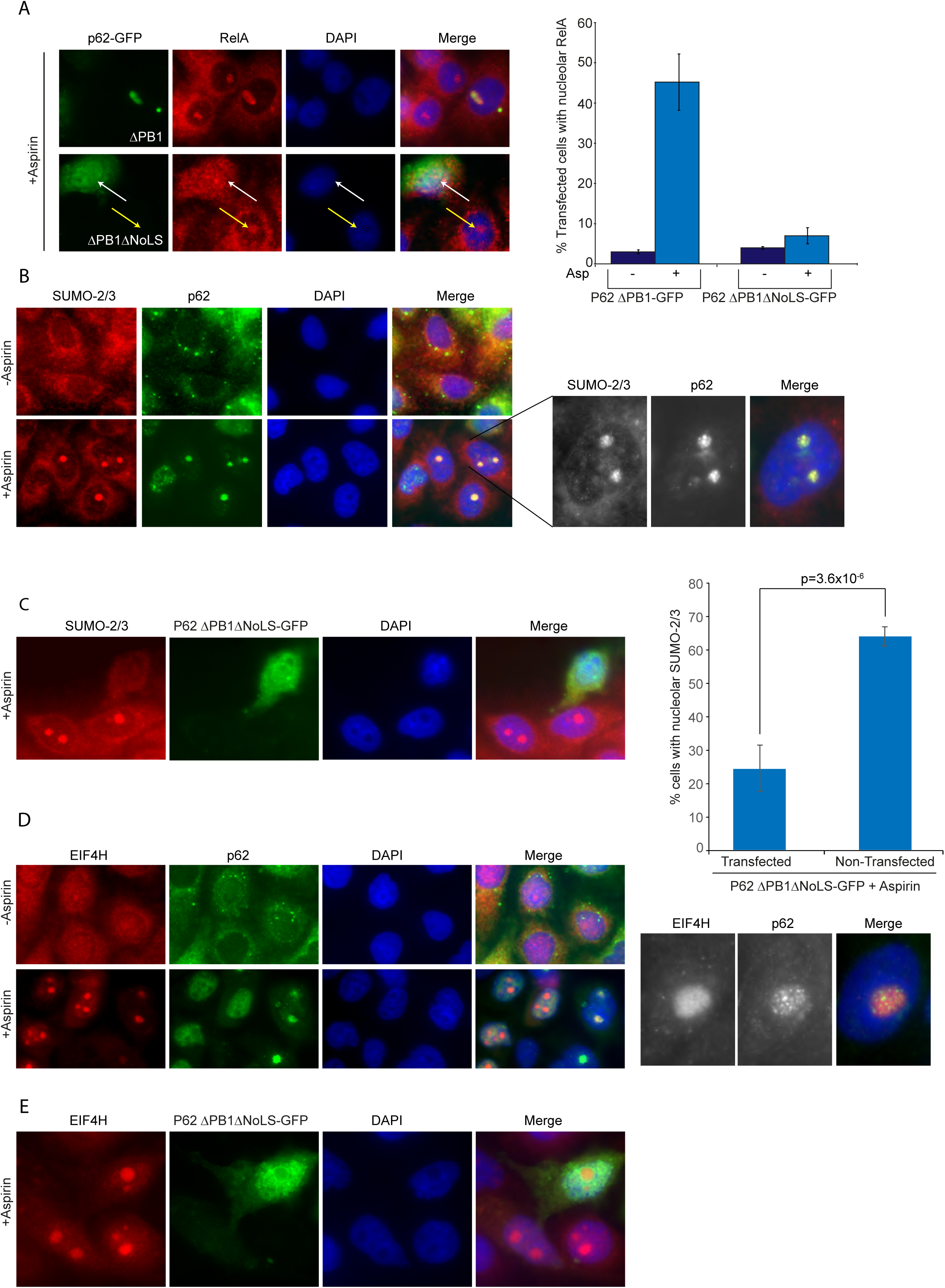
A role for p62 in translocation of RelA to nucleolar aggresomes. **A,C,E** SW480 cells were transfected with GFP-P62ΔPB1 or GFP-p62ΔPBΔNoLS then treated with aspirin (3mM.16h). Immunocytochemistry was performed with RelA (**A**), SUMO2/3 (**B,C**) and EIF4H (D,E) antibodies on fixed cells. **(A)** Right:White arrows indicate transfected cells and yellow arrows indicate nucleolar accumulation of the respective protein in non-transfected cells. The percentage of transfected cells (in a minimum of 100) showing nucleolar RelA (A, left) or nucleolar SUMO2/3 (C, left) was determined. The mean +/− SEM is shown. N=3. P values were derived using a two-tailed students T test. **B and D**, Immunomicrographs (X63) showing the cellular and subcellular localisation of p62 and SUMO2/3/EIF4H with (+) and without (-) aspirin (3mM, 16h) exposure.

To determine whether the reciprocal is true, we utilised depletion of COMMD1 as we have previously shown this very robustly and specifically stops the nucleoplasmic-nucleolar translocation of RelA. Indeed, we found that the aspirin-induced nucleolar accumulation of RelA was blocked upon depletion of COMMD1 (supplemental Fig. 5). However, depletion of COMMD1 had no effect on nucleolar localisation of p62 (Supplemental fig 5), suggesting RelA does not transport p62 to nucleoli.

RelA accumulates in nucleolar aggregates alongside translation factors, chaperones and targets of the ubiquitin proteasome pathway (Fig. 2). Therefore, we considered the role of p62 in nucleolar shuttling of other aggregate proteins. We initially focussed on SUMO-2/3 as this is a classic component of nucleolar aggresomes (26). Immunocytochemistry indicated clear co-localisation between SUMO-2/3 and p62 in nucleolar puncti in response to aspirin (Fig. 7B). Furthermore, expression of GFP-P62ΔPB1ΔNoLS blocked nucleolar translocation of SUMO-2/3 in a similar manner to RelA (Figs 7C). Next, we explored the role of p62 in nucleolar transport of EIF4H, which shows a more diffuse distribution following translocation into the compartment (Fig. 2D). In contrast to RelA and SUMO-2/3, we found little correlation between cells that show nucleolar p62 and those that show nucleolar EIF4H (Fig, 7D). Furthermore, expression of GFP-P62ΔPB1ΔNoLS had little effect on nucleolar translocation of EIF4H (Fig. 7D).

Together, these results reveal that p62 plays a critical role in transporting aggresome-related proteins, including RelA, to nucleoli. They also suggest that this effect is specific and there are p62 dependent and independent mechanisms of nucleolar transport.

## Discussion

The NF-κB transcription factor is one of the most important regulators of the cellular life/death balance and its aberrant activation is associated with many of the hallmarks of cancer. Therefore, identifying means to switch off aberrant NF-κB activity could have a major therapeutic benefit. One such mechanism is nucleolar sequestration of RelA, which causes decreased NF-κB activity and the induction of apoptosis. Indeed, nucleolar sequestration has emerged as a broad mechanism for fine tuning cancer related pathways and maintaining cellular homeostasis. However, understanding of the signals that direct RelA and other extranucleolar proteins to the organelle is limited. Here we report the novel observation that the autophagic cargo receptor, p62, localises to nucleoli. Furthermore, we show that p62 is responsible for transporting specific nuclear proteins, including RelA, to nucleolar aggresomes. These data identify a new role for p62 in the nucleus and shed further light on mechanisms underlying nucleolar sequestration of regulatory proteins that could be used to develop a novel class of therapeutics.

Using fluorescent correlation spectroscopy and quantitative proteomics, we show that aspirin and MG132 cause RelA to accumulate in slow moving nucleolar complexes alongside aggregate-related proteins, suggestive of nucleolar aggresomes. Our conclusion that p62 transports RelA to these aggresomes is based on the following observations. Firstly, we demonstrate that p62 co-localises with RelA and sumoylated proteins in these nucleolar foci. Secondly, we demonstrate that p62 interacts with RelA prior to aggresome formation. We show a correlation between cells that have nucleolar p62 and those that have nucleolar RelA. Finally, we demonstrate that expressing a dominant negative mutant of p62, unable to translocate to nucleoli, specifically blocks nucleolar translocation of RelA. This mutant also blocks nucleolar transport of the aggresome related protein, SUMO2/3, but has no effect on the nucleolar translocation of EIF4H which does not aggregate in nucleoli but is diffuse within the organelle. It will now be of considerable interest to identify the mechanisms by which p62 transports cargo proteins to nucleoli and to understand the determinants of specificity. Given that ubiquitination of RelA is known to be essential for transport of the protein to nucleoli(17), we would predicted that p62 transports ubiquitinated RelA to the organelle via its UBA domain. However, our studies to determine the role of this domain in nucleolar transport of RelA were inconclusive, suggesting it may be partially involved (data not shown).

The mechanisms underlying nucleolar translocation of p62 appear to be distinct from those involved in transport of the protein to autophagosomes. For example, dimerization via the PB1 domain, which is required for autophagasomal translocation, is unnecessary for nucleolar transport(34). It is not enhanced by bafilomycin treatment, which is in keeping with other studies showing nucleolar aggresome formation is unaffected by inhibition of autophagy(41). Furthermore, unlike accumulation at autophagosomes, mutating the LC3 domain has little effect. Indeed, our data would suggest that there is competition between autophagic and nucleolar accumulation of the protein. One domain we did reveal to be essential for translocation of p62 from the nucleoplasm to nucleoli was aa 94-110. This domain is conserved among species and aligns with known NoLSs. We considered that deleting this domain may disturb an essential protein-protein interaction and/or post-translational modification. However, no proteins are documented to bind to this region and prediction programs suggest it is not modified by PTMs. One mechanism suggested for nucleolar accumulation of extra-nucleolar proteins is detention within the organelle via a specific nucleolar detention sequence (NoDS) (42). However, alignment analysis suggests the domain of p62 essential for nucleolar translocation is most likely an NoLS, rather than an NoDS. It will now be interesting to determine what proteins bind to the NoLS of p62 in response to proteotoxic stress and how this signals for nucleolar translocation.

The idea that nucleolar aggresomes act as a means to maintain nuclear proteostasis has gained considerable traction in recent years. Recently, Frottin et al (2019) (43)demonstrated that upon heat stress, misfolded proteins are sequestered in the granular component of nucleoli in liquid phase. Under mild stress, this process is transient and allows misfolded proteins to be maintained in a folding competent state until proteotoxic stress is relieved. However, prolonged stress leads to a transition from liquid-like to solid phase, with loss of reversibility and dysfunction in protein quality control. Others have also suggested that the formation of nucleolar aggresomes can be both transient or irreversible, depending on the stress and context (26). We have previously demonstrated that RelA has a functional impact on proteins within nucleoli that leads to apoptosis (44). Therefore, we suggest that that the RelA-p62 nucleolar aggresomes we observe are not transient, but allow elimination of cells that have acquired irreversible defects in nuclear protein quality control.

Incontrovertible evidence indicates aspirin has anti-tumour properties and the potential to prevent colorectal cancer(45). However, the agent cannot be fully recommended for prevention purpose due to its side effect profile. It is now critical to understand how aspirin acts against cancer cells so that biomarkers of response can be identified. We have previously demonstrated that an early response to aspirin exposure is inhibition of rDNA transcription and the induction of an atypical form of nucleolar stress (29). Here we demonstrate the agent has a particularly significant effect on the nucleolar proteome, compared to the cytoplasmic and nucleoplasmic proteomes. We also show aspirin causes an accumulation of insoluble proteins in colon cancer cells and that these form aggresomes within the organelle. Together, these data suggest that nucleoli play a complex but significant role in the anti-tumour effects of this agent and that nucleolar structure/function could act as a good biomarker of aspirin response.

In summary, we identify a new role for p62 in maintaining nuclear proteostasis by transporting regulatory proteins to nucleolar aggresomes. High levels of nuclear NF-κB activity are a contributory factor in many chronic conditions including cancer. The challenge now is to understand the specifics of proteotoxic stress leading to nucleolar aggregation of p62-RelA, so that we can specifically target RelA to this compartment, thus eliminating diseased cells.

## References

1. DiDonato JA, Mercurio F, Karin M. NF-kappaB and the link between inflammation and cancer. Immunol Rev 2012;246:379–400

2. Perkins ND. The diverse and complex roles of NF-kappaB subunits in cancer. Nat Rev Cancer 2012;12:121–32

3. Gilmore TD. Introduction to NF-kappaB: players, pathways, perspectives. Oncogene 2006;25:6680–4

4. Taniguchi K, Karin M. NF-kappaB, inflammation, immunity and cancer: coming of age. Nat Rev Immunol 2018;18:309–24

5. Prescott JA, Cook SJ. Targeting IKKbeta in Cancer: Challenges and Opportunities for the Therapeutic Utilisation of IKKbeta Inhibitors. Cells 2018;7

6. Concetti J, Wilson CL. NFKB1 and Cancer: Friend or Foe? Cells 2018;7

7. Natoli G. NF-kappaB and chromatin: ten years on the path from basic mechanisms to candidate drugs. Immunol Rev 2012;246:183–92

8. Huang B, Yang XD, Lamb A, Chen LF. Posttranslational modifications of NF-kappaB: another layer of regulation for NF-kappaB signaling pathway. Cell Signal 2010;22:1282–90

9. Chen LF, Greene WC. Shaping the nuclear action of NF-kappaB. Nat Rev Mol Cell Biol 2004;5:392–401

10. Mankan AK, Lawless MW, Gray SG, Kelleher D, McManus R. NF-kappaB regulation: the nuclear response. J Cell Mol Med 2009;13:631–43

11. Campbell KJ, Rocha S, Perkins ND. Active repression of antiapoptotic gene expression by RelA(p65) NF-kappa B. Mol Cell 2004;13:853–65

12. Ankers JM, Awais R, Jones NA, Boyd J, Ryan S, Adamson AD, et al. Dynamic NF-kappaB and E2F interactions control the priority and timing of inflammatory signalling and cell proliferation. Elife 2016;5

13. Stark LA, Dunlop MG. Nucleolar sequestration of RelA (p65) regulates NF-kappaB-driven transcription and apoptosis. Mol Cell Biol 2005;25:5985–6004

14. Loveridge CJ, Macdonald AD, Thoms HC, Dunlop MG, Stark LA. The proapoptotic effects of sulindac, sulindac sulfone and indomethacin are mediated by nucleolar translocation of the RelA(p65) subunit of NF-kappaB. Oncogene 2008;27:2648–55

15. Parrondo R, de las Pozas A, Reiner T, Rai P, Perez-Stable C. NF-kappaB activation enhances cell death by antimitotic drugs in human prostate cancer cells. Mol Cancer 2010;9:182

16. Sniderhan LF, Garcia-Bates TM, Burgart M, Bernstein SH, Phipps RP, Maggirwar SB. Neurotrophin signaling through tropomyosin receptor kinases contributes to survival and proliferation of non-Hodgkin lymphoma. Exp Hematol 2009;37:1295–309

17. Thoms HC, Loveridge CJ, Simpson J, Clipson A, Reinhardt K, Dunlop MG, et al. Nucleolar targeting of RelA(p65) is regulated by COMMD1-dependent ubiquitination. Cancer Res 2010;70:139–49

18. O’Hara A, Simpson J, Morin P, Loveridge CJ, Williams AC, Novo SM, et al. p300-mediated acetylation of COMMD1 regulates its stability, and the ubiquitylation and nucleolar translocation of the RelA NF-kappaB subunit. J Cell Sci 2014;127:3659–65

19. Carlotti F, Chapman R, Dower SK, Qwarnstrom EE. Activation of nuclear factor kappaB in single living cells. Dependence of nuclear translocation and anti-apoptotic function on EGFPRELA concentration. J Biol Chem 1999;274:37941–9

20. Bjorkoy G, Lamark T, Brech A, Outzen H, Perander M, Overvatn A, et al. p62/SQSTM1 forms protein aggregates degraded by autophagy and has a protective effect on huntingtin-induced cell death. J Cell Biol 2005;171:603–14

21. Kavanagh DM, Smyth AM, Martin KJ, Dun A, Brown ER, Gordon S, et al. A molecular toggle after exocytosis sequesters the presynaptic syntaxin1a molecules involved in prior vesicle fusion. Nat Commun 2014;5:5774

22. Ehm P, Nalaskowski MM, Wundenberg T, Jucker M. The tumor suppressor SHIP1 colocalizes in nucleolar cavities with p53 and components of PML nuclear bodies. Nucleus 2015;6:154–64

23. Vilotti S, Codrich M, Dal FM, Pinto M, Ferrer I, Collavin L, et al. Parkinson’s disease DJ-1 L166P alters rRNA biogenesis by exclusion of TTRAP from the nucleolus and sequestration into cytoplasmic aggregates via TRAF6. PLoS One 2012;7:e35051

24. Latonen L, Moore HM, Bai B, Jaamaa S, Laiho M. Proteasome inhibitors induce nucleolar aggregation of proteasome target proteins and polyadenylated RNA by altering ubiquitin availability. Oncogene 2011;30:790–805

25. Audas TE, Jacob MD, Lee S. The nucleolar detention pathway: A cellular strategy for regulating molecular networks. Cell Cycle 2012;11:2059–62

26. Latonen L. Phase-to-Phase With Nucleoli - Stress Responses, Protein Aggregation and Novel Roles of RNA. Front Cell Neurosci 2019;13:151

27. Souquere S, Weil D, Pierron G. Comparative ultrastructure of CRM1-Nucleolar bodies (CNoBs), Intranucleolar bodies (INBs) and hybrid PML/p62 bodies uncovers new facets of nuclear body dynamic and diversity. Nucleus 2015;6:326–38

28. Din FV, Theodoratou E, Farrington SM, Tenesa A, Barnetson RA, Cetnarskyj R, et al. Effect of aspirin and NSAIDs on risk and survival from colorectal cancer. Gut 2010;59:1670–9

29. Chen J, Lobb IT, Morin P, Novo SM, Simpson J, Kennerknecht K, et al. Identification of a novel TIF-IA-NF-kappaB nucleolar stress response pathway. Nucleic Acids Res 2018;46:6188–205

30. Elson EL. Fluorescence correlation spectroscopy: past, present, future. Biophys J 2011;101:2855–70

31. Stark LA, Din FVN, Zwacka RM, Dunlop MG. Aspirin-induced activation of the NF-?B signalling pathway: A novel mechanism for aspirin-mediated apoptosis in colon cancer cells. FASEB J 2001

32. Sanchez-Martin P, Saito T, Komatsu M. p62/SQSTM1 : ‘Jack of all trades’ in health and cancer. FEBS J 2019;286:8–23

33. Lamark T, Svenning S, Johansen T. Regulation of selective autophagy: the p62/SQSTM1 paradigm. Essays Biochem 2017;61:609–24

34. Pankiv S, Lamark T, Bruun JA, Overvatn A, Bjorkoy G, Johansen T. Nucleocytoplasmic shuttling of p62/SQSTM1 and its role in recruitment of nuclear polyubiquitinated proteins to promyelocytic leukemia bodies. J Biol Chem 2010;285:5941–53

35. Feng Y, Duan T, Du Y, Jin S, Wang M, Cui J, et al. LRRC25 Functions as an Inhibitor of NF-kappaB Signaling Pathway by Promoting p65/RelA for Autophagic Degradation. Sci Rep 2017;7:13448

36. Moscat J, Diaz-Meco MT, Wooten MW. Signal integration and diversification through the p62 scaffold protein. Trends Biochem Sci 2007;32:95–100

37. Horke S, Reumann K, Schweizer M, Will H, Heise T. Nuclear trafficking of La protein depends on a newly identified NoLS and the ability to bind RNA. J Biol Chem 2004

38. Emmott E, Dove BK, Howell G, Chappell LA, Reed ML, Boyne JR, et al. Viral nucleolar localisation signals determine dynamic trafficking within the nucleolus. Virology 2008;380:191–202

39. Weber JD, Kuo ML, Bothner B, DiGiammarino EL, Kriwacki RW, Roussel MF, et al. Cooperative signals governing ARF-mdm2 interaction and nucleolar localization of the complex. Mol Cell Biol 2000;20:2517–28

40. Scott MS, Troshin PV, Barton GJ. NoD: a Nucleolar localization sequence detector for eukaryotic and viral proteins. BMC Bioinformatics 2011;12:317

41. Latonen L. Nucleolar aggresomes as counterparts of cytoplasmic aggresomes in proteotoxic stress. Proteasome inhibitors induce nuclear ribonucleoprotein inclusions that accumulate several key factors of neurodegenerative diseases and cancer. Bioessays 2011;33:386–95

42. Audas TE, Jacob MD, Lee S. Immobilization of proteins in the nucleolus by ribosomal intergenic spacer noncoding RNA. Mol Cell 2012;45:147–57

43. Frottin F, Schueder F, Tiwary S, Gupta R, Korner R, Schlichthaerle T, et al. The nucleolus functions as a phase-separated protein quality control compartment. Science 2019;365:342–7

44. Khandelwal N, Simpson J, Taylor G, Rafique S, Whitehouse A, Hiscox J, et al. Nucleolar NF-kappaB/RelA mediates apoptosis by causing cytoplasmic relocalization of nucleophosmin. Cell Death Differ 2011;18:1889–903

45. Chen J, Stark LA. Aspirin Prevention of Colorectal Cancer: Focus on NF-kappaB Signalling and the Nucleolus. Biomedicines 2017;5

