## supplementary figure legends for "A role for the autophagic receptor, SQSTM1/p62, in trafficking NF-κB/RelA to nucleolar aggresomes"

**Supplemental Figure 1: Fluorescence correlation spectroscopy (FCS) curves and fits for DsRed-RelA**

SW480 cells were transfected with DsRed-RelA and GFP-Fibrillarin, treated with aspirin (0-5mM 16h) then analysed by FCS. Average normalised, experimental and fitted autocorrelation curves, with the mean t1 marked (left-hand panels) are shown for DsRed-RelA in the cytoplasm (top), nucleoplasm (middle) and nucleoli (bottom) before and after aspirin treatment. Examples of individual recordings from each compartment are also displayed (right-hand panels).

**Supplemental Figure 2: Aspirin has no significant effect on the diffusion rate of control proteins.** SW480 cells were transfected with pDsRed-C1 and pEGFP-C1 backbone plasmids the treated with aspirin (0-5mM 16h) then analysed by FCS. **(A to C)** Diffusion rate measurements for EGFP (A) and DsRed (B) in each of the three compartments in control and aspirin-treated cells. Combined data indicates there is no significant difference between the two plasmids, or in response to aspirin (C) Error bars represent the SEM and p-values are derived using the Kruskal-Wallis Test. See also Figure 1.

**Supplemental Figure 3 Stable isotope labelling of Amino Acids In Culture (SILAC)-based quantitative proteomics of cellular fractions.**

SW480 cells were cultivated for at least five passages in commercially available DMEM F12 (Dundee Cell products) where arginine and lysine were replaced either by standard amino acids (Arg0; Lys0; light) or by isotope-labelled amino acids (Arg6 and Lys4;medium), or (Arg10 and Lys8; heavy) and supplemented with 10% dialyzed FCS (Invitrogen) and penicillin-streptomycin. Cells were treated with aspirin (6/10h) or MG132 (10h) as indicated. Cells were then mixed and fractionated into cytoplasm, nucleoplasm and nucleoli before being subjected to 1D SDS-PAGE. One biological replicate for 0/6h/10h aspirin and two replicates for 0/MG132/10h aspirin were carried out. This gel was cut into pieces and proteins trypsin digested. The resulting peptides were analyzed by LC-MS/MS and quantified using MaxQuant software. All data was plotted as normalized $Log^2$ values against the control. Negative $Log^2$ values indicate reduced nucleolar abundance whereas positive values indicate increased nucleolar abundance, as compared with the untreated sample. **B** Western blot analysis was performed on isolated cell fractions using antibodies to a-tubulin (cytoplasmic marker), LaminB1 (nucleoplasmic marker) and RPA194 (nucleolar marker). **C** Graph depicting the average number of proteins that increased ($Log^2$ >0.5 and 1) or decreased ($Log^2$<-0.5 and -1) in each compartment (+/- SEM) in response to MG132. The average (Av.) number of proteins isolated from each fraction is given. **D** SW480 cells were treated with aspirin (10mM, hours specified) or MG132 (10uM, 8h) then solubilised using a detergent containing buffer. The concentration of soluble and insoluble proteins were measured then the percentage of insoluble protein determined. Mean (n=3) +/- SEM is shown. * P<0.05 (students T test).

**Supplemental figure 4. Nucleolar p62 and nucleolar RelA are linked at a single cell level.** SW480 cells were treated with aspirin (10mM) for 8hs. Immunomicrographs (63X) show the localisation of p62 and RelA. Arrows indicate cells in which there is no nucleolar p62 or RelA. Scale bar=10um.

**Supplemental Figure 5. Depleting COMMD1 blocks nucleolar translocation of RelA but not, p62**

**A and B** SW480 cells were transfected with control or COMMD1 siRNA **A** Immunomicrographs (X63) showing the localisation of p62 and RelA in the transfected cell population after aspirin (3mM. 16h) exposure. **B** Immunoblot confirming siRNA depletion of COMMD1.
